# Genetic analysis of protein efficiency and its association with performance and meat quality traits under a protein-restricted diet

**DOI:** 10.1101/2022.08.18.503754

**Authors:** Esther Oluwada Ewaoluwagbemiga, Giuseppe Bee, Claudia Kasper

## Abstract

**Background:** An essential component in the development of a sustainable pig production is the reduction of nitrogen excretion in fattening pigs. Pig feeds typically contain high levels of dietary crude protein, and due to incomplete conversion to muscle tissue, excess nitrogen is excreted, resulting in environmental problems such as nitrate pollution and greenhouse gas emissions. Therefore, improving protein efficiency (PE), i.e., the proportion of dietary protein that remains in the carcass, is desirable. This study aimed to estimate the heritability (h^2^) of PE and its genetic correlations with phosphorus efficiency, three performance, seven meat quality and two carcass quality traits when pigs were fed a 20% protein-restricted diet, using a total of 1,071 Swiss Large White pigs. To determine PE, the intake of feed with known nutrient content was accurately recorded for each pig and the nitrogen and phosphorus content of the carcass was determined using dual-energy X-ray absorptiometry.

**Results:** We found an average PE of 0.39 ± 0.04 and a heritability of 0.60 ± 0.08. PE showed a high genetic correlation with phosphorus efficiency (0.68 ± 0.08), moderate genetic correlations with feed conversion ratio (−0.53 ± 0.13) and average daily feed intake (−0.42 ± 0.13), and very little to no genetic correlation with average daily gain (−0.06 ± 0.16). While PE has favourable genetic correlations with the performance traits and some meat quality traits, there is a potentially unfavourable relationship of PE with meat colour (redness [r_g_ = −0.26 ± 0.17]; yellowness [r_g_ = −0.30 ± 0.18]) and intra-muscular fat (IMF; r_g_ = −0.39 ± 0.15). Feed conversion ratio (FCR) also showed unfavourable genetic correlations with meat lightness, redness yellowness, IMF and cooking loss.

**Conclusions:** PE is heritable and can be considered in breeding to reduce the environmental impact of pig production. We found no strong negative influence on meat quality traits (except for meat color and IMF), and there is the potential for indirectly selecting for improved phosphorus efficiency. Selecting nutrient efficiencies might be a more suitable strategy to reduce nitrogen pollution from manure than focusing on FCR because the latter also shows genetic antagonism with some meat quality traits in our population.

## Background

In the past, the focus of animal breeding had been on improving production traits, but today, sustainability concerns are increasingly gaining importance. An essential component to consider in the development of a sustainable pig production chain is the reduction of nitrogen excretion in fattening pigs. Compared to other crops, soybeans contain the highest amount of lysine, which is why soybean meal is often included in the feed to meet the lysine requirements of pigs, resulting in high dietary crude protein content in the feed [1]. Consequently, there is increased excretion of excess nitrogen in feces and urine, which contributes significantly to environmental problems such as nitrate pollution [2] and greenhouse gas emissions in the form of nitrous oxide when manure is applied to pastures and fields [3]. Besides pollution, the large-scale export of soybean from South America has caused massive deforestation in this region, loss of terrestrial biodiversity and deterioration of ecosystem services [4]. In addition, the increasing demand for meat, as a result of rapid growth of the world population, has led to massive expansion of land for soybean cultivation, thereby displacing natural vegetation and other crop cultivation [5]. Lowering the crude protein content in pig feed could help reduce deforestation by lowering the demand for soybean, and therefore make it possible to use locally available protein plant protein sources in countries that import large amounts of soybean meal for pig feed. Therefore, considering the environmental and social impact of pig production, it is essential to improve protein efficiency (PE) in pigs. PE is defined as the proportion of total dietary protein intake that is retained in the carcass. An improvement in PE would simultaneously decrease protein excretion, thereby reducing the contribution of pigs to environmental pollution and greenhouse gas emission. Since more than 50% of the ingested dietary protein in pigs is excreted as waste [6, 7], feed is an important factor to consider in improving PE. Pomar and Remus [9] reported that for every percent reduction in dietary nitrogen, nitrogen excretion could be decreased by 1.5%, thus improving PE. Ruiz-Ascacibar et al. [10], who investigated the influence of a 20% reduction in dietary protein on the PE of the Swiss Large White pig population, reported that pigs fed the 20% protein-reduced diet had significantly longer days on feed in the grower and finisher phase than those on the control diet. For instance, castrated males took 2.5 days longer in both the grower and finisher phase and females took 5.9 and 8.4 days longer in grower and finisher phase, respectively [10]. However, in addition to nutritional strategies to improve PE, a genetic solution can be sought. Ruiz-Ascacibar et al. [9] reported a range of 37.5 - 55% for PE within the protein-reduced treatment group. This variation between individuals to retain dietary protein can be exploited for the purpose of breeding. Additionally, since excreted phosphorus also contributes to environmental pollution, it is interesting to investigate the relationship of PE with phosphorus efficiency for a possible indirect selection of the latter.

Feed conversion ratio (FCR) and residual feed intake (RFI) are the two most common traits in improving feed efficiency, and both are economically relevant. However, ecologically important traits such as PE should also be investigated to achieve a more sustainable pig production while maintaining the same production level. Although FCR and RFI are expected to be correlated with PE, selection for improved FCR and RFI would likely increase energy efficiency rather than PE [6, 7], since energy intake is a main factor driving feed intake [8]. Moreover, in poultry, selection for improved FCR and RFI with the aim of reducing nutrient excretion has been shown to be markedly less efficient than direct selection for the traits nitrogen or phosphorus excretion [11]. Nevertheless, in contrast to FCR and RFI, very few studies have investigated the possibility to improve PE, which may be due to several factors such as difficulties in phenotyping animals for this trait and the lack of approved and validated proxies.

Previous studies using a range of pig breeds, growth phases and diets reported heritabilities of 36-43% for nitrogen retention [12], 21-27% [13] and 36-42% [7] for PE. Nitrogen digestibility coefficient, reflecting the pig’s efficiency to digest dietary fiber and absorb proteins, had an estimated heritability of 27-56% [13]. For total nitrogen excretion during the finisher phase, Shirali et al. [15] reported a heritability (± SE) of 0.32 (± 0.21). Finally, several QTL for nitrogen excretion traits were mapped and their effects and genetic architecture were described [16]. All these studies indicate a potential to select for protein-efficient pigs.

To estimate PE, the amount of protein ingested as well as the amount of protein retained in the animal needs to be determined. While measuring the amount of protein ingested has been greatly facilitated with the use of automatic feeders that record daily feed intake of each pig, determining the amount of protein retention can be laborious, and previously used methods are often either not suitable or too laborious and expensive for large-scale phenotyping. For instance, a direct, but expensive and laborious, method to determine PE is wet-chemistry analysis of the carcass and even the whole body, including blood and organs [7]. Nitrogen retention can be predicted from lean meat content, which is estimated from the weights of primal cuts during dissection [12], which is laborious and subject to variation among butchers [17]. The deuterium dilution technique enables the estimation of nitrogen excretion traits from empty body water content [15] and a nitrogen digestibility coefficient can be estimated from near-infrared spectroscopy of feces [14]. Lean meat content can also be estimated from a range of parameters, such as feed intake and growth patterns [13], or combinations of weight and backfat thickness [18, 19]. These indirect techniques often rely on specific assumptions and might not be generalized to different breeds or sexes, requiring [specific prediction equations for each breed or sex [20]. Indirect methods also yield less precise estimates (R^2^ between 0.94 and 0.990 for predicting chemical values [18], and R^2^ of 0.896 and 0.908 for predicting lean mass from reference dissection [19]), even when specific estimation equations are applied. Here, we use a novel phenotyping strategy including a dual-energy X-ray absorptiometry (DXA) device in combination with automated feeders to estimate PE in a cheaper, more streamlined and faster, but still highly accurate way [21]. Carcass protein content can be estimated by DXA with high precision and accuracy (R^2^ between 0.986 and 0.998 [18]; R^2^ = 0.983 [21]). However, the accuracy for bone mineral content, which we used to compute phosphorus efficiency, is lower [R^2^ 0.886 and 0.875 for empty body and carcass, respectively [21]; R^2^ between 0.816 and 0.851 [39], for predicting ash from DXA bone mineral content). DXA has been successfully applied to genetic studies of body composition in pigs [22] and other livestock species (reviewed in [20]). Moreover, in contrast to estimation methods that rely on point measurements, such as backfat thickness or loin muscle area, the information of total body as well as region-specific composition provided by DXA and other whole-body scan methods enables the targeted improvement of specific areas of the carcass through breeding programs.

The knowledge of genetic correlations of PE with other traits of importance (such as meat quality and performance traits) is also important to account for possible trade-offs with traits included in current breeding programs. Studies have reported both favourable and unfavourable genetic relationships between the commonly used feed efficiency traits (FCR and RFI) with meat quality traits and average daily gain [12, 23]. Phosphorus efficiency and its genetic correlation with PE is also an important trait to consider, as the low N:P ratio in pig feces results in excess phosphorus in soils when pig manure is used as a fertilizer [24, 25]. In addition, it is necessary to assess whether, at least on a phenotypic basis, PE could be associated with customers’ acceptance of meat based on flavor, tenderness and juiciness.

The major aim of this study was therefore to estimate heritability of PE and its genetic correlations with phosphorus efficiency, production and meat quality traits when dietary protein is reduced. In addition, we aim to estimate heritability of performance traits (ADG, ADFI and FCR) and meat quality traits, and their genetic correlations. Finally, we assessed differences in the sensory evaluation of meat between different PE groups based on tenderness, flavor and juiciness.

## Methods

### Animals and data sets

A total of 1,071 pigs using combined datasets from previous experiments (294 pigs [7]; 48 pigs [26]; 48 pigs in a further study on protein and essential amino-acid reduction in the growth and finisher period or solely in the finisher period [27]; and an additional 681 pigs raised specifically for this study) were used in the analysis of PE (Table 1). In all experiments explained below, pigs had *ad libitum* access to isocaloric diets that differed in crude protein or fibre content. Furthermore, in all experiments, pigs were fed a grower diet from 20 to 60 kg body weight and a finisher diet from 60 to slaughter at 100 kg. In some experiments, pigs were kept until a target weight of 140 kg and fed another specially formulated finisher diet from 100 to 140 kg [10]. In all experiments, control diets were formulated according to the Swiss feeding recommendations for pigs^1^. The data from previous experiments at Agroscope [28] included four experimental runs with eight farrowing series (i.e. batches of litters born in the same week), and pigs were assigned either to the control or protein-restricted diets. The protein-restricted diets contained 80% of the crude protein and digestible essential amino acids content of the respective control diets. The data collected by Bee et al. [26] included two experimental runs and pigs were assigned to three experimental treatments (T95, T100, and T100-CF). Pigs in the T95 treatment were fed the control diets that complied with the BIOSUISSE regulation, which allows 95% of the feed ingredients being of organic origin^2^. The diets in the T100 treatment consisted of 100% feed ingredients that complied with the aforementioned regulation. The diet used in the T100-CF treatment was the same formulation as the T100, but with the crude fibre content increased to 6% by including sunflower press cake, sainfoin and lupine. Pigs in this experiment were slaughtered at a target BW of 105 kg. The 48 pigs from Bee et al. [27] were assigned to three experimental treatments: C, the control diet that contained 80% of the crude protein and digestible essential amino acids content of the respective control diet only in the finisher, but with the control diet in the grower phase (RPF), and another one with a protein reduction in both, grower and finisher phase (RPGF). Due to similarities between treatment groups in all experiments described above, treatment groups were pooled over different experiments (Table 1). Pigs were slaughtered at a target body weight of 100 kg. Finally, the 681 pigs that were raised and phenotyped specifically for this study, originated from 14 farrowing series (39 sires and 79 dams in total), and data were collected from October 2018 to June 2021. Forty-eight dams had one litter, 23 dams had two litters and eight dams had three litters. All sires and dams were of the Swiss Large White breed. All pigs in this experiment were fed a RPGF diet. Pigs in this experiment were slaughtered at an average body weight of 106 ± 5 kg. The target weight of 100 kg for slaughter was chosen to reflect current Swiss remuneration standards for carcasses, which penalize slaughter weights below 80 and above 100 kg.

**Table 1:**
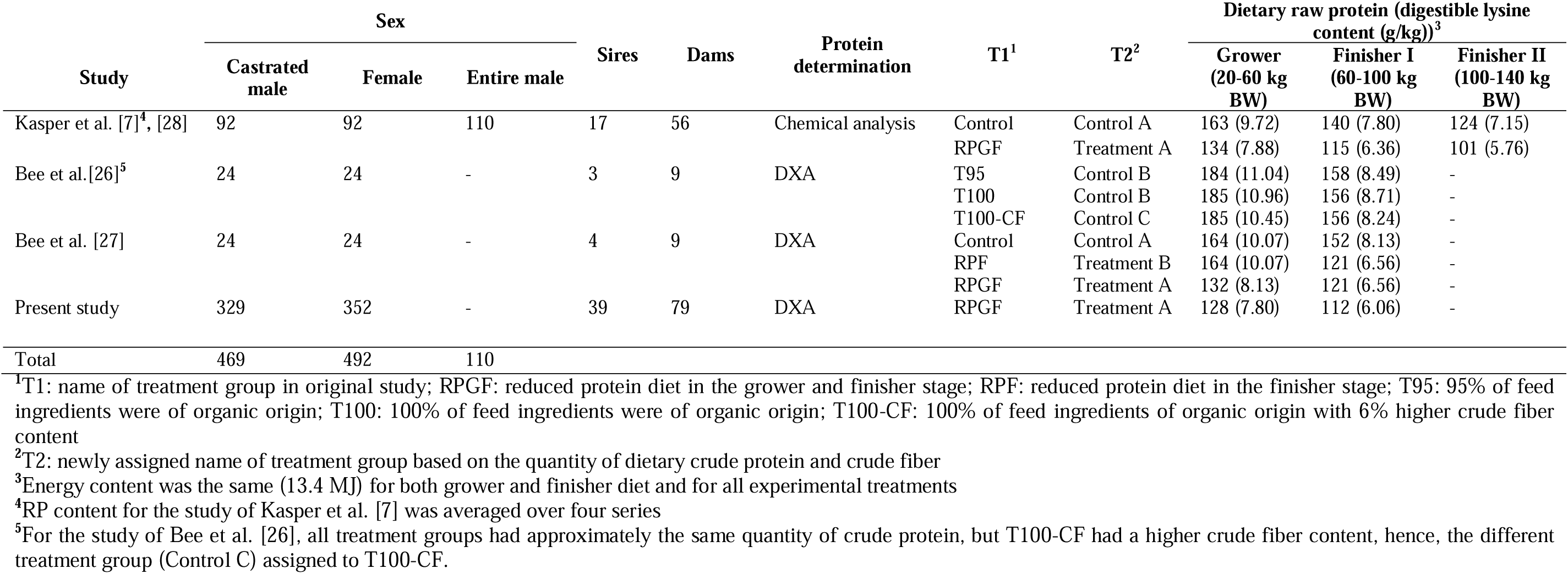
Overview of datasets and the origin of the 1,071 pigs used in the study, indicating the differences in the number of animals, sexes, and methods to determine carcass protein content, treatment groups, dietary crude protein and digestible lysine content.

Piglets were weaned at an average age of 27 ± 2 days after birth by removing the sow and were fed a standard starter diet with crude protein levels following the recommendation. At 22.3 (± 1.6) kg, pigs were placed in pens equipped with automatic feeders (single-spaced automatic feeder stations with individual pig recognition system by Schauer Maschinenfabrik GmbH & Co. KG, Prambachkirchen, Austria) and stayed on the starter diet. The experiment started at this stage and all pigs learned to access the automatic feeders, which allows monitoring of feed intake. The automatic feeder recorded all feeder visits and feed consumption per visit, from which the total feed intake of each pig was calculated [29]. The protein content of feed was monitored during production by near-infrared spectroscopy for each 500 kg batch. Each week, a sample was taken from each automatic feeder station and the crude protein content was determined by wet-chemistry methods. This was done to adjust for fluctuations in the crude protein content of raw materials when calculating PE, as the diet was formulated at the start of the study based on tabulated values of the ingredients. Every week, pigs were weighed individually, and, once the pig reached the target start weight of 20 kg, they were allocated to grower-finisher pens and the experimental treatments were started. This was done until a maximum number of 12 (or 24 or 48) pigs per pen (depending on the pen layout; minimum 1m^2^ per pig and maximum 12 pigs/feeder) was reached. Pigs remained in this pen until slaughter.

Total and average daily feed (ADFI) was recorded, and average daily gain (ADG) and the feed conversion ratio (FCR) were calculated as;

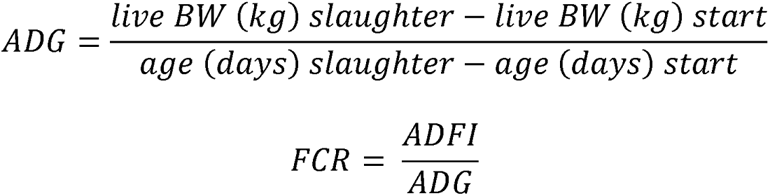

where *live BW* (*kg*) *slaughter* and *age* (*days*) *slaughter* are the live pre-slaughter body weight in kg and the age in days at slaughter, respectively, and and *live BW* (*kg*) *start* and *age* (*days*) *start* are the exact body weight in kg and the age in days at the start of the grower phase, respectively.

### Protein and phosphorus efficiency

For the data in Kasper et al. [7], pigs were serially slaughtered at 20 – 140 kg body weight in 20 kg intervals and the nitrogen and phosphorus content of the carcasses was determined by wet chemical analysis following the protocol described by [10]. For the other data sets, pigs were slaughtered at an average body weight of 106 ± 5 kg and the left carcass half, including the whole head and the tail, was scanned with a Dual-energy x-ray absorptiometry (DXA; GE Lunar i-DXA, GE Medical Systems, Glattbrugg, Switzerland) to determine the lean tissue and bone mineral content. The lean tissue content and bone mass obtained from the DXA scans was used in the following prediction equation to estimate the protein content retained in the carcass [21].

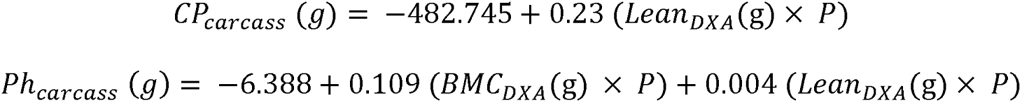

Where *CP_carcass_* (*g*) is the crude protein content of the carcass in g, *Lean_DXA_* (*g*) is the lean meat content obtained with DXA in g, *P* is the proportion of the weight of the left cold carcass-half weight (including the whole head and the tail) to the total cold carcass weight, *Ph_carcass_* (*g*) is the phosphorus content of the carcass in g, and *BMC_DXA_* (g) is the bone mineral content obtained with DXA in g. Protein and phosphorus efficiency of the carcass was thereafter calculated as the ratio of protein (or phosphorus) retained in the carcass (corrected for protein (or phosphorus) content in the carcass at a target live body weight of 20 kg, which is the start of the experiment and when feed intake was first monitored) to the total protein (or phosphorus) intake (*CP_feed_ intake* (g)) during the experimental period.

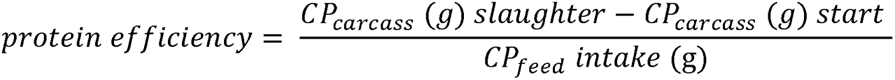

The protein and phosphorus content of pigs at the start of this experiment *CP_carcass_* (*g*) *start* was estimated from a sample of 38 piglets (12 females, 12 castrated males and 14 entire males). These 38 piglets were slaughtered at an average of 20.98 ± 1.85 kg body weight in a previous experiment and their carcass protein content was chemically determined [10]. The average protein content per kg carcass for each sex (female, entire male, castrated male) was used to estimate *CP_carcass_* (*g*) *start* for pigs by multiplying the actual live body weight of pigs when they entered the experiment (i.e., at approximately 20 kg body weight) with the protein content per kg carcass of piglet, as previously determined from the 38 piglets [10].

### Meat quality traits and predictor variables in the study

In order to investigate possible trade-offs between PE and other traits of importance, such as meat quality and performance, additional traits were recorded on a subset of pigs raised specifically for this study (N=509; Table 2). After exsanguination and evisceration, backfat thickness was measured at the 10^th^ rib level on the hot left carcass side with a ruler. Thereafter, the eviscerated carcasses (left and right half) were weighed and then stored overnight at 4 °C. One day after slaughter, the longissimus thoracii (LT) muscle was excised (at the 10^th^ to 12^th^ rib level) from the left cold carcass side. The area of the LT was measured at the 11^th^ to 12^th^ rib level. A 1x pixel JPEG image of the muscle was taken with a smartphone camera (always mounted on the same support structure to guarantee the same angle and distance to the object) together with a ruler for scale. Subsequently, the area was measured with the imageJ software (v1.53r). In addition, a 3-cm thick chop, labeled A and B, was cut. After a 20 min bloom period at 4°C, L* (lightness), a* (redness), and b* (yellowness) values for chop A were measured using a spectrophotometer (model CM-2600d, Minolta, Dietikon, Switzerland). Subsequently, from the same chop, drip loss was assessed as the quantity of purge generated during storage at 4 °C for 48 h. Thereafter, the samples were vacuum-sealed in plastic bags and cooked in a water bath at 72 ^°^C for 45 min, then cooled in cold water for 15 min, rinsed to remove coagulated sarcoplasmic protein, dabbed dry and weighed to determine the cooking loss. The cooked chops were then stored at −20 ^°^C until measuring the shear force. Shear force was then determined on the cooked samples, which were slowly brought to ambient temperature. We measured shear force from five cores of each LT chop with a diameter of 1.27 cm perpendicular to the fiber direction using the Stable Micro System TA.XT2 Texture Analyzer (Godalming, Surry, UK) equipped with a 2.5-mm-thick Warner-Bratzler shear. The chops labeled B were freed from subcutaneous fat and then used for the determination of the intramuscular fat content. The samples were placed in plastic bags, vacuumed-sealed and frozen at −20 °C until analysis. The frozen samples were lyophilized using a Delta 2-24 machine (Christ Delta 2-24, Kühner AG, Birsfelden, Switzerland) and the intramuscular fat content was determined by extracting with petrol ether after an acid hydrolysis (International Organization for standardization (ISO), 1999). Drip loss and cooking loss were calculated as;

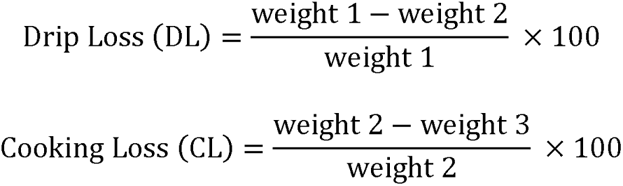

**Table 2:** Traits measured in addition to protein and phosphorus efficiency and predictor variables used in analysis

| **Variable category** | **Trait/variable** | **Unit** |
| --- | --- | --- |
| Nutrient efficiency<br>(N=1,071) | Protein efficiency | proportion |
|  | Phosphorus efficiency | proportion |
| Meat quality<br>(N=510) | Meat colour (lightness, redness and yellowness) | International Commission on Illumination (CIE) color space coordinates |
|  | Intra-muscular fat content | g/kg |
|  | Water holding capacity (drip loss, cooking loss) | % |
|  | Shear force | N |
|  | Backfat thickness | mm |
|  | Loin muscle area | cm<sup>2</sup> |
| Sensory evaluation of meat<br>(N= 39) | Firmness | Rated by judges on a linear gradual scale ranging from 0.0 to 10.0 |
|  | Tenderness |  |
|  | Juiciness |  |
|  | Flavour |  |
| Performance traits<br>(N=1,071) | Average daily gain (ADG) | kg/d |
|  | Feed conversion rate (FCR) | kg/kg |
|  | Average daily feed intake (ADFI) | kg/d |
| Predictor variables<br>(N=1,071) | Temperature | °C |
|  | Sex | - |
|  | Treatment | - |
|  | Year of change | Calendar year |
|  | Slaughter weight | kg |
|  | Slaughter age (rAgeLW; residuals of slaughter age on slaughter weight) | days |

**Table 3:** Description of variables used in the models for efficiency, performance and meat quality traits

| Variable | Included in model |  | Levels | Variable type |
| --- | --- | --- | --- | --- |
|  | Univariate | bivariate |  |  |
| Year of change | Yes | Yes | 1 | Fixed continuous |
| Slaughter weight (kg) | Yes | Yes | 1 | Fixed continuous |
| Slaughter age (days) | Yes | Yes | 1 | Fixed continuous |
| Temperature (°C) | Yes | Yes | 1 | Fixed continuous |
| Treatment<sup>1</sup> | Yes | Yes | 5 | Fixed factor |
| Sex | Yes | Yes | 3 | Fixed factor |
| Animal | Yes | Yes | 1071 | Random factor |
| Sibship<sup>2</sup> | Yes | Yes | 197 | Random factor |
| Interactions<sup>3</sup> | Yes | - |  |  |
| Treatment:slaughter age |  |  |  |  |
| Year:slaughter age |  |  |  |  |
| Slaughter weight:sex |  |  |  |  |
| Treatment:sex |  |  |  |  |

<sup>1</sup>The variable ‘Treatment’ was used only in a univariate and bivariate model of efficiency and performance traits (PE, phosphorus efficiency, ADG, FCR, ADFI)
<sup>2</sup>The variable ‘Sibship’ was used only in a univariate and bivariate model of efficiency and performance traits (PE, phosphorus efficiency, ADG, FCR, ADFI)
<sup>3</sup>Interactions for meat quality traits differed depending on the top model selected for each meat quality trait.

**Table 4:** Description of variables used in the model for the sensory analysis

| Variable | Values | Number | Variable type |
| --- | --- | --- | --- |
| Judge |  | 8 | Random factor |
| Session |  | 6 | Random factor |
| Sex |  |  | Fixed factor |
|  | Castrated males | 18 |  |
|  | Females | 21 |  |
| Protein efficiency |  |  | Fixed factor |
|  | HPE | 12 |  |
|  | LPE | 14 |  |
|  | MPE | 13 |  |
| Fat thickness |  |  | Fixed factor |
|  | <1.5 cm | 18 |  |
|  | 1.5 cm | 12 |  |
|  | 3 cm | 9 |  |

HPE: high protein-efficient; LPE: low protein-efficient; MPE: medium protein-efficient

Where weight 1 is the weight of the chop after a 20 mins bloom period at 4°C, weight 2 is the weight after 48h at 4°C, and weight 3 is the weight after cooking in a water bath at 72 ^°^C for 45 min and cooled in cold water for 15 min with coagulated sarcoplasmic protein removed and dabbed dry. Predictor variables such as slaughter weight, slaughter age, treatment, year of change, sex and temperature were included in the genetic analysis of the efficiency traits, performance traits and meat quality traits. Slaughter weight is the weight of the animal at the time of slaughter and slaughter age is the age of the animal at slaughter. Due to high correlation of slaughter age with slaughter weight (Pearson r = 0.77, *p* < 0.001), we used the residuals of a linear regression of slaughter age on slaughter weight (rAgeLW) in place of slaughter age to avoid collinearity. Year of change is the year the animal entered into the experiment with an average body weight of 20 kg. Temperature is the average ambient temperature (± 3 days) in the room at the time the animal entered the experiment.

### Sensory Evaluation of Meat

The objective of the sensory evaluation was to identify potential differences in the perceived initial firmness (first bite), tenderness, juiciness and flavour of meat samples between three groups differing in PE (low, LPE; medium, MPE; and high, HPE). Chops of 2 cm thickness were selected from pigs raised specifically for this study. Pigs with a PE less than the mean PE - 2σ were considered as LPE (group average PE of 0.35 ± 0.08), pigs with a PE greater than mean PE + 2σ were considered as HPE (group average PE of 0.57± 0.05), and pigs in between were considered as MPE group (group average PE of 0.40 ± 0.01). A total number of 12 chops (from 6 castrates and 6 females), 13 chops (from 6 castrates and 7 females), and 14 chops (from 6 castrates and 8 females) were used in HPE, MPE, and LPE, respectively. Chops were taken between the 11^th^ and 12^th^ ribs (the same chop used in measuring loin muscle area) of the left cold carcass half 24h postmortem and stored at −80 ^°^C. For the sensory analysis, chops were thawed for 24h at 4°C, blotted dry and stored at room temperature for approximately 1h. Slices were grilled at 180 °C for 2.45 min on each side and cut into pieces of approximately 2 × 2 cm. The chops differed in the thickness of the attached fat (<1.5 cm, 1.5 cm or 3 cm), which we corrected statistically in the model. The cut samples were kept at 60°C until sensory testing. A panel of eight judges participated in the sensory tests. The intensity of the attributes firmness, tenderness, juiciness, and total flavour was measured on an unstructured, gradual 10-cm line scale between low (0) and high (10) intensity. Prior to data collection, panelists participated in two training sessions to get familiar with the attributes of interest. In each of the six sessions, panelists evaluated two sets, each of them consisting of meat cuts of three different animals. Sample sets and samples within each test set were randomized according to a William Latin Square design. All samples were coded with three-digit random numbers. Sensory data were collected using the software FIZZ (version 2.51 Biosystèmes, Couternon, France). White bread, still water and warm black tea were provided for neutralization of the mouth between samples. All tests were conducted at room temperature under daylight conditions in the sensory laboratory at Agroscope Posieux.

### Statistical analysis

All analyses were performed in R software V 4.2.1 [30]. The *pedantics* package V 1.7 [31] was used to construct a multigenerational pedigree for the animal model and to derive pedigree-based parameters such as relatedness and inbreeding. The total pedigree contained 1468 unique individuals with a maximum pedigree depth of 12 generations. From these individuals, 682 pigs were pigs raised specifically for this study, 390 pigs were from previous experiments (described in Kasper et al. [7], as well as Bee et al. [26] and Bee et al. [27]), and 396 pigs with no phenotypes provided links for the individuals with PE data. All phenotyped individuals had both parents known. There was a mean inbreeding coefficient of 0.00032. The genetic analysis (heritabilities and correlations) was performed using the variance and covariances obtained with *ASReml-R* [32]. All analyses were also done in *MCMCglmm* [33] for general comparison, and in particular to make them comparable to a previous study [7] (Supplementary Information Figures S1 – S2, Table S1). To determine the effect of several variables on PE, we first ran a linear model that included all those variables and interactions as fixed effects. Interactions were included to account for the heterogeneity of data, i.e., in some experiments, entire males were used whereas in the others only females and castrated males were present, and some experiments had different dietary groups (Table 1). The structure of our data set was not fully cross-classified. Thus, as the model results of these interactions are not of interest per se, we do not discuss them in this work. We then selected the top model(s), i.e., all models within delta AIC < 2 with the ‘dredge’ function in *MuMin* package V 1.46.0 [34]. The variables retained in the top model were then used as fixed effects in the univariate animal model to estimate the genetic and common environmental variance components. The univariate animal models were performed using the model formula;

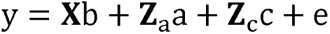

Where ***y*** is a vector of observations of the respective trait, ***b*** is a vector of fixed effects of the year the pig entered the experiment (year of change), slaughter weight, residuals of slaughter age on slaughter weight (rAgeLW), experimental treatment, sex, temperature, treatment × rAgeLW interaction, slaughter weight × sex interaction, treatment × sex interaction, and year of change × rAgeLW interaction. **X** is an incidence matrix relating records to fixed effects, *a* is a vector of random additive genetic effects, and **Z** is the corresponding incidence matrix. *c* is a vector of random litter effects, **Z*_c_*** is the corresponding incidence matrix, and *e* is the random residual effect. Heritability was thereafter computed as the ratio of genetic variance to the phenotypic variance (*h^2^* = *V_A_* /*V_P_*), where phenotypic variance (*V_P_*) is the sum of genetic variance (*V_A_*), litter (common environmental) effect (*V_CE_*) and residual variance (*V_R_*). The litter effect was calculated as the ratio of litter/common environment variance to phenotypic variance (*CE^2^*= *V_CE_*/*V_P_*). The model distributions were visualized using the plot() method in ASReml-R.

Genetic correlations were estimated using bivariate models including the additive genetic and common environment covariance (the latter only for the correlations among the performance and efficiency traits), in which we added year, slaughter weight, treatment, sex, temperature, slaughter age (rAgeLW) as fixed effects. An unconstrained variance/covariance matrix was assumed for the random models. For bivariate analyes that included litter effect, we specified Starting values by using the argument start.values = TRUE, which allows the model to change its random parameters. Genetic correlations were computed by rescaling additive genetic covariances by the variances.

While all data (N=1071) were used in the heritability estimation of PE and performance traits and their genetic correlations, heritability estimation of meat quality was done on a subset (N=510), for which these phenotypes were available. The univariate animal models used in the heritability estimation of meat quality traits did not contain the fixed effect of treatment since the subset, for which meat quality measurements were available, had only one treatment group. The interactions between covariates identified in the model selection step differed between meat quality traits depending on the top model for each trait. A mixed-effects model in XLSTAT v 2021 was used for the phenotypic analysis of the sensory evaluation of meat to correct for the effect of session and judge, and Fisher’s LSD mean separation test were performed for post-hoc analysis. The model used was;

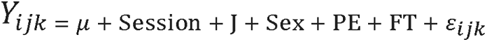

Where *Y_ijk_* are the observations for the dependent variables (initial firmness, tenderness, juiciness and flavour), is the overall mean, *Session* is the random effect for session, *μ* is the random effect for judge, *Sex* is the fixed effect of sex (castrates and females), *PE* is the fixed effect of PE group (HPE, LPE and MPE), and *FT* is the fixed effect of the thickness of fat adhering to chops.

## Results

### Influence of fixed effects on PE, phosphorus efficiency, FCR, ADG and ADFI

The descriptive statistics for the efficiency traits, FCR (kg/kg), ADG (kg/day), and ADFI (kg/day) are presented in Table 5 and those for meat and carcass quality, as well as for the sensory evaluation of meat, are presented in Table 6. Using the entire dataset (1,071 pigs), PE, phosphorus efficiency, FCR, ADG and ADFI had a mean of 0.39 ± 0.04, 0.43 ± 0.05, 2.67 ± 0.23, 0.85 ± 0.11, and 2.26 ± 0.31, respectively (Table 5). Slaughter weight, experimental treatments, sex, age (rAgeLW), and ambient temperature showed significant influence on PE (P < 0.05; Table 7). PE of pigs decreased by 0.7%, 0.4% and 1.2% as the slaughter weight (P = 0.001), ambient temperature (P = 0.020) and age (rAgeLW) (P = <0.001) increased, respectively. While pigs in Treatment B (i.e., pigs fed a reduced protein diet at the finishing stage) were not statistically different from pigs in control A (i.e., pigs fed the standard diet) (P = 0.276), pigs in Treatment A, Control B, and Control C were statistically different from pigs in Control A (P = <0.001). Pigs in Treatment A (i.e., pigs fed a reduced protein diet in the grower and finisher period) had 1.8% higher PE compared to pigs in control A (P = <0.001). Pigs in control B (i.e., pigs fed the organic diet with RP content higher than the control A) and control C (i.e., pigs fed the organic diet and crude fibre with RP content higher than the control A) had 5.1% and 6.5% lower PE compared to those in control A (P = <0.001), respectively. Sex showed significant influence on PE, with females having 1.3% higher PE than the castrated males (P = 0.007) and entire males having 1.7% higher PE than the castrated males (P = <0.001).

**Table 5:** Descriptive statistics of traits for each study and the overall dataset

| Trait | Study<sup>1</sup> | N | Mean | SD |
| --- | --- | --- | --- | --- |
| Protein efficiency | Bee et al. [26] | 48 | 0.34 | 0.04 |
|  | Bee et al. [27] | 48 | 0.39 | 0.04 |
|  | Present study | 681 | 0.40 | 0.03 |
|  | **Overall<sup>2</sup>** | 1071 | 0.39 | 0.04 |
| Phosphorus efficiency | Bee et al. [26] | 48 | 0.37 | 0.04 |
|  | Bee et al. [27] | 48 | 0.40 | 0.03 |
|  | Present study | 681 | 0.42 | 0.03 |
|  | **Overall<sup>2</sup>** | 1071 | 0.43 | 0.05 |
| FCR | Bee et al. [26] | 48 | 2.54 | 0.15 |
|  | Bee et al. [27] | 48 | 2.59 | 0.11 |
|  | Present study | 681 | 2.76 | 0.14 |
|  | **Overall<sup>2</sup>** | 1071 | 2.67 | 0.23 |
| ADG | Bee et al. [26] | 48 | 0.95 | 0.06 |
|  | Bee et al. [27] | 48 | 0.94 | 0.07 |
|  | Present study | 681 | 0.84 | 0.11 |
|  | **Overall<sup>2</sup>** | 1071 | 0.85 | 0.11 |
| ADFI | Bee et al. [26] | 48 | 2.40 | 0.19 |
|  | Bee et al. [27] | 48 | 2.44 | 0.18 |
|  | Present study | 681 | 2.32 | 0.26 |
|  | **Overall<sup>2</sup>** | 1071 | 2.26 | 0.31 |

<sup>1</sup>Descriptive statistics for the data of Kasper et al. [28] is reported in the study of Kasper et al. [7]
<sup>2</sup>Descriptive statistics for “**Overall**” also includes the data from Kasper et al. [7]

**Table 6:** Descriptive statistics of traits for meat quality traits and meat sensory evaluation of meat protein efficiency groups

|  | Trait | N | Mean | SD |
| --- | --- | --- | --- | --- |
| Meat quality | IMF | 509 | 44.57 | 15.58 |
|  | LMA | 509 | 35.35 | 3.80 |
|  | L* | 509 | 51.82 | 2.28 |
|  | a* | 509 | 6.55 | 0.98 |
|  | b* | 509 | 3.91 | 0.89 |
|  | Shear force | 509 | 44.94 | 11.09 |
|  | Drip loss | 509 | 2.45 | 0.97 |
|  | Cooking loss | 509 | 26.91 | 2.67 |
|  | Backfat thickness | 509 | 21.17 | 5.60 |
| Sensory evaluation | HPE | 12 | 0.57 | 0.05 |
|  | MPE | 13 | 0.40 | 0.01 |
|  | LPE | 14 | 0.35 | 0.08 |

**Table 7:** Influence of fixed effects on efficiency traits

| Variable | Protein efficiency |  | Phosphorus efficiency |  |
| --- | --- | --- | --- | --- |
|  | Estimate ± SE | P<sup>1</sup> | Estimate ± SE | P<sup>1</sup> |
| **Intercept** | 0.373 ± 0.005 | <0.001 | 0.436 ± 0.007 | <0.001 |
| **Year** | 0.003 ± 0.003 | 0.250 | -0.001 ± 0.004 | 0.66 |
| **Slaughter weight** | -0.007 ± 0.002 | 0.001 | -0.015 ± 0.003 | <0.001 |
| **Treatment<sup>2</sup>** |  | <0.001 |  | <0.001 |
| Treatment B | 0.019 ± 0.017 | 0.276 | -0.035 ± 0.027 | 0.184 |
| Treatment A | 0.018 ± 0.005 | <0.001 | -0.018 ± 0.007 | 0.011 |
| Control B | -0.051 ± 0.014 | <0.001 | -0.068 ± 0.023 | 0.003 |
| Control C | -0.065 ± 0.015 | <0.001 | -0.084 ± 0.024 | <0.001 |
| **Sex<sup>3</sup>** |  | <0.001 |  | <0.001 |
| Females | 0.013 ± 0.005 | 0.016 | -0.000 ± 0.008 | 0.968 |
| Males | 0.017 ± 0.006 | 0.007 | 0.023 ± 0.009 | 0.015 |
| **Slaughter age (rAgeLW)** | -0.012 ± 0.002 | <0.001 | -0.015 ± 0.004 | <0.001 |
| **Temperature** | -0.004 ± 0.002 | 0.020 | -0.008 ± 0.002 | 0.002 |
| **Treatment:slaughter age** |  | <0.001 |  | <0.001 |
| Treatment B:slaughterAge | 0.009 ± 0.013 | 0.471 | 0.021 ± 0.021 | 0.298 |
| Treatment A:slaughterAge | 0.018 ± 0.002 | <0.001 | 0.017 ± 0.003 | <0.001 |
| ControlB:slaughterAge | 0.011 ± 0.011 | 0.358 | 0.018 ± 0.018 | 0.335 |
| ControlC:slaughterAge | 0.045 ± 0.02 | 0.009 | 0.040 ± 0.027 | 0.144 |
| **Year:slaughter age** | -0.002 ± 0.001 | 0.031 | -0.006 ± 0.002 | <0.001 |
| **Slaughter weight:sex** |  | <0.001 |  | 0.002 |
| Slaughter weight:female | 0.002 ± 0.002 | 0.430 | 0.002 ± 0.003 | 0.608 |
| Slaughter weight:male | 0.016 ± 0.002 | <0.001 | 0.013 ± 0.004 | 0.001 |
| **Treatment:sex** |  | 0.002 |  | 0.204 |
| Treatment B:Female | 0.002 ± 0.015 | 0.898 | 0.031 ± 0.024 | 0.189 |
| Treatment A:Female | -0.003 ± 0.006 | 0.586 | 0.019 ± 0.009 | 0.029 |
| Treatment A:Male | -0.022 ± 0.007 | 0.001 | 0.000 ± 0.011 | 0.973 |
| Control B:Female | 0.026 ± 0.01 | 0.018 | 0.032 ± 0.017 | 0.059 |
| Control C:Female | -0.012 ± 0.014 | 0.405 | 0.010 ± 0.022 | 0.647 |

<sup>1</sup>P-values are converted from z-scores provided by ASReml-R<sup>2</sup>Control A was included in the intercept<sup>3</sup>Castrated male (castrates) was included in the intercept

For phosphorus efficiency, there was a significant influence of slaughter weight, ambient temperature, and age (rAgeLW), with pigs having 1.5% (P < 0.05), 0.8% (P < 0.05) and 1.5% (P < 0.05) lower phosphorus efficiency as slaughter weight, ambient temperature and age (rAgeLW) increased, respectively (Table 7). Experimental treatments showed statistical significant influence on phosphorus efficiency (P < 0.05), with treatment A, control B, and control C having 1.8% (P < 0.05), 6.8% (P < 0.05) and 8.4% (P < 0.05) lower phosphorus efficiency compared to control A, respectively (Table 7). Sex also showed significant influence on phosphorus efficiency, with entire males having 2.4% higher PE compared to castrated males (P = 0.015) (Table 7).

Table 8 shows the influence of the fixed effects on the performance traits. For FCR, there was a significant influence of slaughter weight, experimental treatments, sex and age (rAgeLW) (P < 0.05). Pigs had 17% increased FCR as the slaughter weight increases (P = <0.001) and 9% lower FCR as the age (rAgeLW) increased (P = <0.001). Pigs in Treatment A and Control C had 17% (P = <0.001) and 16% (P = 0.03) increased FCR compared to pigs in Control A. Females and entire males had 7% (P = 0.005) and 10% (P = 0.001) lower FCR than the castrated males. For ADG and ADFI, there was a significant influence of year of change, slaughter weight, experimental treatments, sex and age (rAgeLW). ADG of pigs decreased by 2% as the slaughter weight increased (P < 0.05) and ADG increased by 9% as age (rAgeLW) increased (P < 0.05). Only pigs in Treatment A showed significant difference from pigs in control A for ADG (P = <0.001), with pigs in Treatment A having 3% lower ADG compared to the pigs in control A. Castrated pigs were significantly different from females and entire males with females and entire males having lower ADG than the castrated males (P < 0.05). Finally, ADFI increased as the slaughter weight and age (rAgeLW) increased (P < 0.05). Only pigs in Treatment A showed a significant difference from pigs in control A for ADFI (P = 0.003), with pigs in Treatment A having higher ADFI compared to the pigs in control A. Castrated pigs were significantly different from females and entire males with females and entire males having lower ADFI compared to the castrated males (P = <0.001).

**Table 8:** Influence of fixed effects on performance traits

|  |  | FCR |  | ADG |  | ADFI |  |
| --- | --- | --- | --- | --- | --- | --- | --- |
| Variable |  | Estimate ± SE | P<sup>1</sup> | Estimate ± SE | P<sup>1</sup> | Estimate ± SE | P<sup>1</sup> |
| Intercept |  | 2.57 ± 0.02 | <0.001 | 0.89 ± 0.01 | <0.001 | 2.28 ± 0.02 | <0.001 |
| Year |  | 0.02 ± 0.01 | 0.111 | 0.01 ± 0.00 | 0.008 | 0.04 ± 0.01 | 0.001 |
| Slaughter weight |  | 0.17 ± 0.01 | <0.001 | -0.02 ± 0.00 | <0.001 | 0.08 ± 0.01 | <0.001 |
| Treatment<sup>1</sup> |  |  | <0.001 |  | 0.000 |  | <0.001 |
|  | Treatment B | 0.11 ± 0.08 | 0.215 | -0.02 ± 0.03 | 0.596 | 0.07 ± 0.08 | 0.411 |
|  | Treatment A | 0.17 ± 0.02 | <0.001 | -0.03 ± 0.01 | <0.001 | 0.06 ± 0.02 | 0.003 |
|  | Control B | 0.04 ± 0.07 | 0.597 | -0.01 ± 0.03 | 0.726 | 0.03 ± 0.07 | 0.628 |
|  | Control C | 0.16 ± 0.08 | 0.034 | -0.03 ± 0.00 | 0.317 | 0.10 ± 0.07 | 0.180 |
| Sex<sup>2</sup> |  |  | <0.001 |  | <0.001 |  | <0.001 |
|  | Females | -0.07 ± 0.03 | 0.005 | -0.03 ± 0.01 | <0.001 | -0.15 ± 0.02 | <0.001 |
|  | Males | -0.10 ± 0.03 | 0.001 | -0.03 ± 0.01 | 0.002 | -0.18 ± 0.03 | <0.001 |
| Slaughter age (rAgeLW) |  | -0.09 ± 0.01 | <0.001 | 0.09 ± 0.00 | <0.001 | 0.18 ± 0.01 | <0.001 |
| Temperature |  | 0.00 ± 0.01 | 0.707 | 0.01 ± 0.00 | 0.087 | 0.02 ± 0.01 | 0.047 |
| Treatment:slaughter age |  |  | 0.001 |  | 0.004 |  | 0.954 |
|  | TreatmentB:slaughterAge | -0.04 ± 0.06 | 0.521 | 0.01 ± 0.02 | 0.631 | -0.03 ± 0.06 | 0.671 |
|  | TreatmentA:slaughterAge | -0.05 ± 0.01 | <0.001 | 0.02 ± 0.00 | <0.001 | 0.00 ± 0.01 | 0.757 |
|  | ControlB:slaughterAge | -0.01 ± 0.06 | 0.908 | 0.02 ± 0.02 | 0.401 | 0.01 ± 0.06 | 0.803 |
|  | ControlC:slaughterAge | 0.05 ± 0.08 | 0.521 | 0.01 ± 0.03 | 0.778 | 0.05 ± 0.08 | 0.543 |
| Year:slaughter age |  | -0.01 ± 0.01 | 0.136 | 0.00 ± 0.00 | 0.631 | 0.00 ± 0.01 | 0.683 |
| Slaughter weight:sex |  |  | 0.003 |  | 0.157 |  | 0.038 |
|  | Slaughter weight:female | -0.02 ± 0.01 | 0.031 | 0.00 ± 0.00 | 0.199 | -0.01 ± 0.01 | 0.171 |
|  | Slaughter weight:male | -0.04 ± 0.01 | 0.001 | 0.01 ± 0.00 | 0.064 | -0.03 ± 0.01 | 0.011 |
| Treatment:sex |  |  | <0.001 |  | 0.168 |  | 0.125 |
|  | Treatment B:Female | 0.02 ± 0.07 | 0.847 | 0.02 ± 0.02 | 0.485 | 0.06 ± 0.07 | 0.379 |
|  | Treatment A:Female | 0.02 ± 0.03 | 0.591 | 0.00 ± 0.01 | 0.776 | 0.01 ± 0.03 | 0.600 |
|  | Treatment A:Male | -0.12 ± 0.03 | 0.001 | 0.03 ± 0.01 | 0.010 | -0.03 ± 0.03 | 0.347 |
|  | Control B:Female | -0.08 ± 0.05 | 0.129 | 0.00 ± 0.02 | 0.867 | -0.08 ± 0.05 | 0.124 |
|  | Control C:Female | -0.09 ± 0.07 | 0.205 | 0.00 ± 0.02 | 0.971 | -0.08 ± 0.07 | 0.208 |

<sup>1</sup>P-values are converted from z-scores provided by ASReml-R<sup>2</sup>Control A was included in the intercept<sup>3</sup>Castrated male (castrates) was included in the intercept
Abbreviations: FCR (feed conversion ratio); ADG (average daily gain); ADFI (average daily feed intake)

### Heritability and litter effects

Table 9 summarizes the estimates of heritabilities and variance components of PE and several performance and meat quality traits of Swiss Large White pigs. The result showed a heritability estimate ± SE of 0.60 ± 0.08 for PE in the carcass. The contribution of the litter effect to the phenotypic variance in carcasses for PE was approximately zero (5.8 × 10^-8^ ± 0), which showed that growing up in the same early environment did not lead to those pigs’ PE being more similar beyond the additive genetic effect. The heritability estimates of phosphorus efficiency, ADG, FCR, and ADFI were 0.46 ± 0.11, 0.59 ± 0.11, 0.47 ± 0.10, and 0.59 ± 0.11, respectively (Table 9). In addition, there was a litter effect of 0.038 ± 0.037, 0.10 ± 0.04, 0.039 ± 0.035 and 0.08 ± 0.04 to phosphorus efficiency, ADG, FCR and ADFI, respectively, with ADG and ADFI having more influence of the early common environment. The heritability estimates of meat quality and carcass traits were moderate to high ranging from 0.34 ± 0.11 in the LT area to 0.73 ± 0.12 in intramuscular fat (Table 9). In general, heritabilities could be estimated with high levels of confidence, as reflected in the low standard error (Table 9).

**Table 9:** Estimates ± SE of additive genetic variance, heritability, common environment variance, litter effect and phenotypic variance of nutrient efficiency, performance, meat quality and carcass traits and their sample sizes.

| Trait | N | $\sigma^2A$ | Heritability | $\sigma^2CE$ | Litter effect | $\sigma^2P$ |
| --- | --- | --- | --- | --- | --- | --- |
| PE | 1071 | $6.1 \times 10^{-4} \pm 1.1 \times 10^{-4}^{***}$ | $0.60 \pm 0.08$ | $5.8 \times 10^{-8} \pm NA$ | $5.7 \times 10^{-5} \pm 3.3 \times 10^{-6}$ | $1.0 \times 10^{-3} \pm 5.9 \times 10^{-5}$ |
| PhE | 1071 | $1.1 \times 10^{-3} \pm 2.9 \times 10^{-4}^{***}$ | $0.46 \pm 0.11$ | $9.1 \times 10^{-5} \pm 8.8 \times 10^{-5}$ | $0.04 \pm 0.037$ | $2.4 \times 10^{-3} \pm 1.3 \times 10^{-4}$ |
| ADG | 1071 | $1.9 \times 10^{-3} \pm 4.3 \times 10^{-4}^{***}$ | $0.59 \pm 0.11$ | $3.2 \times 10^{-4} \pm 1.3 \times 10^{-4}^*$ | $0.10 \pm 0.04$ | $3.2 \times 10^{-3} \pm 2.1 \times 10^{-4}$ |
| FCR | 1071 | $1.1 \times 10^{-2} \pm 2.7 \times 10^{-3}^{***}$ | $0.47 \pm 0.10$ | $9.2 \times 10^{-4} \pm 8.3 \times 10^{-4}$ | $0.04 \pm 0.035$ | $2.4 \times 10^{-2} \pm 1.3 \times 10^{-3}$ |
| ADFI | 1071 | $1.5 \times 10^{-2} \pm 3.4 \times 10^{-3}^{***}$ | $0.59 \pm 0.11$ | $1.9 \times 10^{-3} \pm 9.6 \times 10^{-4}^*$ | $0.08 \pm 0.04$ | $2.5 \times 10^{-2} \pm 1.6 \times 10^{-3}$ |
| L* | 509 | $1.67 \pm 0.53^{**}$ | $0.37 \pm 0.10$ | - | - | $4.47 \pm 0.33$ |
| a* | 509 | $0.49 \pm 0.12^{***}$ | $0.57 \pm 0.11$ | - | - | $0.86 \pm 0.07$ |
| b* | 509 | $0.27 \pm 0.08^{***}$ | $0.41 \pm 0.10$ | - | - | $0.65 \pm 0.05$ |
| LMA | 509 | $4.53 \pm 1.75^{**}$ | $0.34 \pm 0.11$ | - | - | $13.33 \pm 1.13$ |
| IMF | 509 | $144.70 \pm 33.65^{***}$ | $0.73 \pm 0.12$ | - | - | $199.30 \pm 17.96$ |
| ShF | 509 | $39.56 \pm 11.90^{**}$ | $0.42 \pm 0.10$ | - | - | $94.94 \pm 7.15$ |
| DL | 509 | $0.59 \pm 0.17^{***}$ | $0.39 \pm 0.09$ | - | - | $1.52 \pm 0.11$ |
| CL | 509 | $5.90 \pm 1.83^{**}$ | $0.36 \pm 0.09$ | - | - | $16.22 \pm 1.17$ |

\*\*\* = $P < 0.001$ , \*\* = $P < 0.01$ , \* = $P < 0.05$
Abbreviations: PE, protein efficiency; PhE, phosphorus efficiency; ADG, average daily gain; FCR, feed conversion ratio; ADFI, average daily feed intake; L\*, meat lightness; a\*, meat redness; b\*, meat yellowness; LMA, loin muscle area; IMF, intramuscular fat; ShF, shear force; DL, drip loss; CL, cooking loss; BT, backfat thickness. N, sample size, $\sigma^2A$ , additive genetic variance; $\sigma^2CE$ , common environment variance; $\sigma^2P$ , phenotypic variance.

### Genetic and phenotypic correlations

#### Efficiency and performance traits

As PE is a new efficiency trait that is closer to the goal of sustainability, we sought to assess potential trade-offs with the traits commonly included in breeding goals, such as FCR and other performance traits. The genetic correlations between PE and performance traits were moderate and clearly different from zero. PE showed significant genetic correlations with phosphorus efficiency (0.68 ± 0.08), ADFI (−0.42 ± 0.13), and FCR (−0.53 ± 0.13), and a low genetic correlation with ADG (−0.06 ± 0.16) (Table 10). The phenotypic correlations of PE with phosphorus efficiency (0.55 ± 0.03) and performance traits, ADFI (−0.34 ± 0.04), FCR (−0.50 ± 0.03) and ADG (0.07 ± 0.04), had similar patterns with the genetic correlations (Table 10). While the relationship of PE with phosphorus efficiency, ADFI and FCR is favourable (i.e., pigs with higher PE would be phosphorus efficient, consume less feed and efficiently convert the feed), the relationship between PE and ADG does not change (i.e., an increase in PE would neither decrease nor increase daily growth). FCR showed favourable genetic correlations of −0.31 ± 0.15 and 0.44 ± 0.13 with ADG and ADFI, respectively (Table 10). Similarly, phenotypic correlations of FCR with ADG and ADFI were also favourable.

**Table 10:**
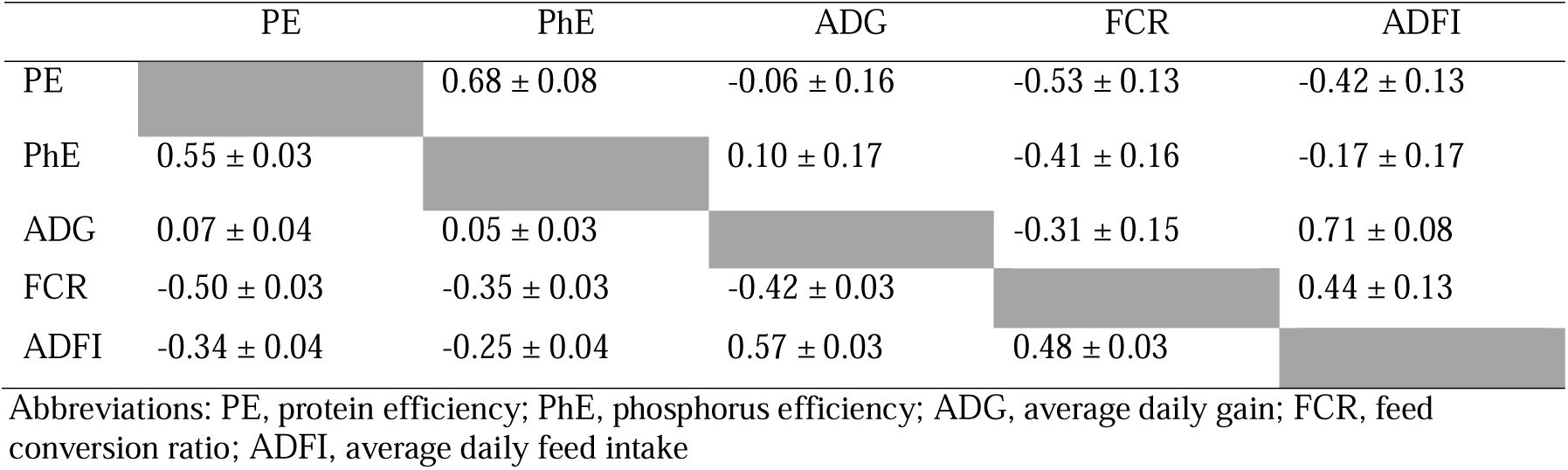
Estimates of genetic correlations (above diagonals) and phenotypic correlations (below diagonals) between protein efficiency and performance traits.

|  | PE | PhE | ADG | FCR | ADFI |
| --- | --- | --- | --- | --- | --- |
| PE | | $0.68 \pm 0.08$ | $-0.06 \pm 0.16$ | $-0.53 \pm 0.13$ | $-0.42 \pm 0.13$ |
| PhE | $0.55 \pm 0.03$ | | $0.10 \pm 0.17$ | $-0.41 \pm 0.16$ | $-0.17 \pm 0.17$ |
| ADG | $0.07 \pm 0.04$ | $0.05 \pm 0.03$ | | $-0.31 \pm 0.15$ | $0.71 \pm 0.08$ |
| FCR | $-0.50 \pm 0.03$ | $-0.35 \pm 0.03$ | $-0.42 \pm 0.03$ | | $0.44 \pm 0.13$ |
| ADFI | $-0.34 \pm 0.04$ | $-0.25 \pm 0.04$ | $0.57 \pm 0.03$ | $0.48 \pm 0.03$ | |

Abbreviations: PE, protein efficiency; PhE, phosphorus efficiency; ADG, average daily gain; FCR, feed conversion ratio; ADFI, average daily feed intake

#### Efficiency, meat quality and carcass traits

PE shows favourable genetic correlations with LMA (0.72 ± 0.16) and backfat thickness (−0.37 ± 0.12) (Table 11). However, PE may have potentially unfavourable genetic correlations with meat redness (a*), meat yellowness (b*), IMF and cooking loss, though the standard error for the correlations are quite high compared to the (Table 11) estimates. A similar pattern was also observed with the phenotypic correlations of PE with meat quality traits (Table 11). FCR showed unfavourable genetic correlations with meat colour (L* = −0.25 ± 0.18, a* = 0.49 ± 0.15, and b* = 0.40 ± 0.17), IMF (0.31 ± 0.16) and cooking loss (−0.46 ± 0.17) (Table 12). The same pattern holds for the phenotypic correlations of FCR with meat colour, IMF and cooking loss.

**Table 11:** Estimates of genetic correlations (above diagonals) and phenotypic correlations (below diagonals) between protein efficiency and meat quality traits.

|  | PE | L* | a* | b* | LMA | IMF | ShF | DL | CL | BT |
| --- | --- | --- | --- | --- | --- | --- | --- | --- | --- | --- |
| PE |  | 0.06 ± 0.19 | -0.26 ± 0.17 | -0.30 ± 0.18 | 0.72 ± 0.16 | -0.39 ± 0.15 | 0.006 ± 0.19 | 0.23 ± 0.19 | 0.59 ± 0.15 | -0.37 ± 0.12 |
| L* | 0.02 ± 0.05 |  | -0.08 ± 0.19 | 0.34 ± 0.17 | 0.31 ± 0.24 | 0.29 ± 0.17 | -0.61 ± 0.13 | 0.12 ± 0.21 | 0.17 ± 0.21 | 0.06 ± 0.20 |
| a* | -0.11 ± 0.06 | 0.17 ± 0.05 |  | 0.85 ± 0.06 | -0.36 ± 0.20 | 0.43 ± 0.14 | -0.46 ± 0.15 | -0.13 ± 0.19 | -0.49 ± 0.16 | -0.23 ± 0.17 |
| b* | -0.11 ± 0.05 | 0.55 ± 0.04 | 0.77 ± 0.02 |  | -0.32 ± 0.22 | 0.59 ± 0.13 | -0.67 ± 0.11 | -0.14 ± 0.20 | -0.40 ± 0.18 | -0.07 ± 0.11 |
| LMA | 0.22 ± 0.06 | 0.04 ± 0.06 | -0.19 ± 0.06 | -0.12 ± 0.06 |  | -0.04 ± 0.21 | -0.02 ± 0.25 | 0.51 ± 0.21 | 0.93 ± 0.11 | 0.14 ± 0.23 |
| IMF | -0.19 ± 0.06 | 0.27 ± 0.05 | 0.26 ± 0.06 | 0.35 ± 0.05 | -0.14 ± 0.06 |  | -0.24 ± 0.17 | -0.31 ± 0.17 | -0.19 ± 0.18 | 0.14 ± 0.17 |
| ShF | 0.02 ± 0.05 | -0.59 ± 0.03 | -0.36 ± 0.05 | -0.53 ± 0.04 | 0.04 ± 0.06 | -0.20 ± 0.06 |  | -0.02 ± 0.20 | 0.08 ± 0.20 | 0.08 ± 0.19 |
| DL | 0.14 ± 0.06 | 0.03 ± 0.05 | -0.02 ± 0.05 | -0.04 ± 0.05 | 0.24 ± 0.05 | -0.16 ± 0.05 | 0.02 ± 0.05 |  | 0.38 ± 0.18 | -0.11 ± 0.20 |
| CL | 0.17 ± 0.05 | -0.08 ± 0.05 | -0.19 ± 0.05 | -0.16 ± 0.05 | 0.43 ± 0.05 | -0.17 ± 0.05 | 0.20 ± 0.05 | 0.39 ± 0.04 |  | 0.23 ± 0.19 |
| BT | -0.18 ± 0.05 | 0.07 ± 0.05 | 0.03 ± 0.06 | 0.08 ± 0.05 | 0.06 ± 0.06 | 0.12 ± 0.06 | -0.09 ± 0.05 | -0.01 ± 0.05 | -0.04 ± 0.05 |  |

Abbreviations: PE, protein efficiency; L\*, meat lightness; a\*, meat redness; b\*, meat yellowness; LMA, loin muscle area; IMF, intramuscular fat; ShF, shear force; DL, drip loss; CL, cooking loss; BT, backfat thickness.

**Table 12:** Estimates of genetic correlations and phenotypic correlations between performance traits and meat quality and carcass traits with their 95% credible intervals in parentheses. Estimates, whose credible intervals do not span zero are printed in bold.

| Genetic correlations |  |  |  |  |  |  |  |  |  |
| --- | --- | --- | --- | --- | --- | --- | --- | --- | --- |
|  | **L\*** | **a\*** | **b\*** | **LMA** | **IMF** | **ShF** | **DL** | **CL** | **BT** |
| **ADG** | 0.12 ± 0.19 | 0.15 ± 0.17 | 0.23 ± 0.17 | -0.17 ± 0.22 | 0.14 ± 0.16 | -0.19 ± 0.18 | -0.01 ± 0.19 | -0.03 ± 0.19 | 0.31 ± 0.16 |
| **FCR** | -0.25 ± 0.18 | 0.49 ± 0.15 | 0.40 ± 0.17 | -0.68 ± 0.18 | 0.31 ± 0.16 | -0.11 ± 0.19 | -0.08 ± 0.19 | -0.46 ± 0.17 | 0.21 ± 0.17 |
| **ADFI** | -0.03 ± 0.18 | 0.37 ± 0.14 | 0.39 ± 0.15 | -0.53 ± 0.17 | 0.29 ± 0.15 | -0.20 ± 0.17 | -0.02 ± 0.18 | -0.28 ± 0.17 | 0.40 ± 0.14 |
| Phenotypic correlations |  |  |  |  |  |  |  |  |  |
|  | **L\*** | **a\*** | **b\*** | **LMA** | **IMF** | **ShF** | **DL** | **CL** | **BT** |
| **ADG** | 0.06 ± 0.06 | 0.14 ± 0.06 | 0.14 ± 0.06 | -0.11 ± 0.06 | 0.17 ± 0.07 | -0.08 ± 0.06 | -0.001 ± 0.06 | -0.08 ± 0.06 | 0.25 ± 0.06 |
| **FCR** | -0.13 ± 0.05 | 0.12 ± 0.06 | 0.05 ± 0.05 | -0.15 ± 0.06 | 0.11 ± 0.06 | -0.002 ± 0.05 | -0.20 ± 0.05 | -0.16 ± 0.05 | 0.10 ± 0.05 |
| **ADFI** | -0.01 ± 0.06 | 0.20 ± 0.06 | 0.17 ± 0.06 | -0.21 ± 0.06 | 0.25 ± 0.06 | -0.09 ± 0.06 | -0.12 ± 0.06 | -0.19 ± 0.05 | 0.31 ± 0.05 |
| Abbreviations: ADG, average daily gain; FCR, feed conversion ratio; ADFI, average daily feed intake; L*, meat lightness; a*, meat redness; b*, meat yellowness; LMA, loin muscle area; IMF, intramuscular fat; ShF, shear force; DL, drip loss; CL, cooking loss; BT, backfat thickness |  |  |  |  |  |  |  |  |  |

### Sensory evaluation of meat

The results showed significant influence of judge and session on all the sensory attributes (P < 0.01), except for the influence of session on juiciness. A significant influence of PE group on juiciness could be observed (P < 0.01). However, MPE and LPE were not significantly different for juiciness, but a significant difference was observed between LPE and HPE as well as between MPE and HPE. This likely means that judges graded the juiciness of LPE and MPE meat samples equally, while they graded HPE samples as less juicy (P < 0.05). PE group did not show significant influence on initial firmness, tenderness and flavor (P > 0.05), suggesting that an improvement in PE may not influence the sensory attributes considered in this study except for juiciness. However, our results should be followed-up with a consumer panel to investigate whether consumers perceive the meat of the distinct PE groups as different.

## Discussion

In this study, we estimated the heritability of PE, phosphorus efficiency and a range of performance, meat and carcass quality traits and their genetic correlations in a population of Swiss Large White pigs under various levels of dietary protein and amino acid supply. As already suggested previously [7], we now clearly show a major potential of selecting protein efficient pigs, mostly without any apparent expected negative impact on production, meat and carcass quality traits. The analysis of environmental influence shows that PE is also affected by ambient temperature, protein content of the feed, sex and age. Therefore, in addition to selection, there are various management practices that can be used to improve PE.

### Protein and phosphorus efficiency

Similar to the study of Kasper et al. [7] and Ruiz-Ascacibar et al. [10], this study also found an effect of treatment group (diet), sex and slaughter weight on protein and phosphorus efficiency. The heritability estimate reported in this study was higher than that reported previously for the same population [7], which may be due to the higher sample size of the current study, in which the previous data was integrated. It is interesting to note that the heritability estimates using only the 681 pigs raised specifically for this study, which were fed reduced-protein diet in the growing and finishing phase, were similar to the overall result (0.62 ± 0.11 vs. 0.60 ± 0.08). Although only few studies have investigated PE in pigs, some have probed similar nutrient efficiency traits, such as total nitrogen excretion and nitrogen digestibility coefficient, and have reported varying heritabilities depending on the breed, growth phase, and diet [11, 12, 13, 14].

Our study showed a higher heritability for PE compared to most of the previous studies. These differences could be due to differences in breed and diet, but also to differences in the methods of nutrient efficiency estimation, which, in our case, might be more reliable compared to the use of indirect estimation methods, such as the deuterium technique or the estimation from lean meat content from dissections or production parameters. Whole-body scans might yield higher heritability estimates of body composition and thus PE than deriving body composition from point measurements such as backfat thickness by ultrasound, due to the reduction of error. However, the difference in the magnitude of the heritability estimate to Kasper et al. [7], where carcass protein was determined by wet chemistry, a very accurate method, is more likely a result of the different statistical models (MCMCglmm) used. Applying the model of Kasper et al. [7] to the data of this study resulted in a highly similar heritability estimate 0.43 [0.29, 0.58] (Supplementary Information Figure S1).

Consistent with previous work, the present study confirms that nutrient efficiency traits are heritable and can be harnessed to substantially reduce environmental pollution in pig production. However, in the practical implementation of breeding, it should be kept in mind that PE as a ratio trait may have a disadvantage. As argued by Zetouni et al. [35], direct selection for a ratio trait such as PE may not lead to the desired result, as it is not certain whether the improvement comes from the counter trait (e.g., protein retention), the denominator trait (e.g., protein intake), or both. Therefore, it is advisable to use a multi-trait selection approach (i.e., selecting for both protein retention and protein intake) to achieve the highest genetic gain for a ratio trait [35]. In addition to direct genetic effects, the composition of the gut microbiome has been shown to influence general feed efficiency [36] and the genetic make-up of the host together with the composition of the microbiome explains more of the phenotypic variation in digestive efficiency of nitrogen than additive genetic variation alone [14]. Thus, this important source of phenotypic variation should potentially be considered in breeding programs.

### Performance traits

This study reports an influence of slaughter weight, sex and treatment group on average daily gain (ADG), feed conversion ratio (FCR) and average daily feed intake (ADFI), which is similar to the results of Kasper et al. [7] and Ruiz-Ascacibar et al. [10]. The heritabilities for performance traits were also mostly in line with those of other studies. We found moderate heritabilities for ADG and ADFI, with the one of ADG being similar to that reported by Shirali et al. [15] of 0.64 ± 0.19. However, they were higher than the estimates reported by Verschuren [13] (h^2^ = 0.27 and h^2^ = 0.43 for ADG and ADFI, respectively) and Kavlak and Uimari [37] (h^2^ = 0.25 ± 0.06 for ADG). FCR was also clearly heritable in the present study (h^2^ = 0.47 ± 0.10), which is similar to, although slightly higher than, the estimates reported by Saintilan et al. [12]. In contrast, Shirali et al. [15] reported a lower heritability of 0.26 ± 0.20 over a period of 60 – 140 kg body weight. Previously, we also reported very low heritability estimates for FCR and ADG [7]. These differing estimates compared to the present study could be due to the higher sample size in the present study, in which there were an additional 777 individuals available. Heritability for FCR may differ depending on growth phase and test period (e.g., at only the grower or finisher phase or throughout all growth phases) as reported by some studies where multiple QTLs for ADG and for FCR at different growth phases [16, 38, 39]. Our moderate heritability estimates for meat quality traits are similar to those of previous studies [15, 40, 41, 42]. As in our study, the redness of the meat consistently showed higher heritability than the lightness and yellowness of the meat. Comparing the estimates with those obtained in MCMCglmm, we find that they are similar, except for IMF, where the estimate was higher in MCMCglmm (Supplementary Information Table S1). The heritability estimate of 0.99 (0.58 – 0.99) for IMF from MCMCglmm seems unrealistically high, given that the estimates in the literature are moderate (h^2^=0.52±0.13 [40] and (h^2^=0.43±0.06 [41]). The bimodal posterior distribution (Supplementary Information Figure S2) suggests that it may be a prior effect that could cause this bias, and taking the posterior mean leads to an overestimation of heritability in this particular case. The first mode of the posterior distribution (Supplementary Information Figure S2) is in fact close to the estimate from ASReml-R (h^2^=0.73±0.12).

### Genetic and phenotypic correlations with performance traits

Favourable genetic correlations were observed between PE and phosphorus efficiency as well as FCR and ADFI. This result shows that selection for increased PE would result in pigs that are also more phosphorus-efficient, consume less feed and efficiently convert the feed into body (and, in particular, muscle) mass. We observed very little to no relationship between PE and ADG, showing that selection for increased PE would likely not influence the daily growth of pigs, though a negative but statistically not significant genetic correlation was observed. Similar relationships were found in the work of Déru et al. [14] on nitrogen digestible coefficient and of Verschuren [13] on PE. In the latter study, the genetic relationships of PE with ADG, FCR and ADFI became stronger with the pigs’ age. This suggests that the genetic factors underlying these traits and their covariances may vary across growth phases. Considering the ecological footprint and feed costs, a focus on the finishing phase would therefore be more relevant, as most feed is consumed in this period. A possible unfavourable relationship between PE and ADG may be due to an indirect effect of lower feed intake since there is a negative relationship between PE and ADFI. However, the relationship between PE and ADG may not be linear, i.e., protein-efficient pigs do not necessarily grow slower during each growth phase. For example, Verschuren [13] reported a low positive, i.e., favourable, genetic relationship (r_G_ = 0.11) between PE and ADG in the starter phase, but a low negative (r_G_ = −0.11) in the grower phase as well as a stronger, unfavourable relationship (r_G_ = −0.43) in the finisher phase. Furthermore, Shirali et al. [16] also found QTLs for nitrogen excretion to be unfavourably associated with ADG only during the 90 kg to 120 kg body weight growth phase. Saintilan et al. [12] also reported favourable genetic correlations of nitrogen excretion efficiency with ADG, FCR and ADFI, but the one with FCR was close to 1, in contrast to r_G_ = −0.53 ± 0.13 in our study. The reason for some of the discrepancy observed between this study and that of Saintilan et al. [12] thus may be due to differences in methodology, traits, and sample size, but most importantly probably due to the mixing of growth phases.

### Genetic and phenotypic correlations with meat and carcass quality traits

In this study, while PE does not show significant correlations with meat lightness (L*) and shear force (i.e., these traits are most likely not influenced by PE), we found favourable genetic correlations of PE with LMA and backfat thickness (i.e., an increase in PE will increase loin muscle area and decrease backfat thickness). Considering the high genetic correlation of PE with LMA, the latter could be considered in predicting PE, but requires the slaughter of the animal. The intermediate negative correlation of BT with PE renders BT rather less interesting as proxy for PE, as a considerable loss of genetic progress in PE would have to be expected. The same applies to dressing percentage, which also had a rather low genetic correlation with PE (rG = 0.28 ± 0.19; Supplementary Information Table S2). PE showed an unfavourable genetic correlation with meat colour (redness and yellowness), IMF and cooking loss. However, the genetic correlations of PE with meat redness and yellowness were moderate, but clearly different from zero. This indicates that genetic improvement of PE might reduce the redness and yellowness of the meat (the meat might look paler), which could be perceived as unattractive by consumers. However, this may depend on the starting value of the meat colour. Déru et al. [14] reported non-significant genetic correlations of nitrogen digestible coefficient with meat lightness, redness and yellowness, which were in an unfavourable direction for meat lightness and redness depending on the type of diet (conventional or high fiber diet). Additional experiments, such as the visual assessment and preferences of meat colour by consumers, should be conducted to investigate whether differences in meat colour within the expected range would lead to different consumer decisions. The sensory analysis in our study suggests no apparent conflicts of PE with the way the judges perceived the initial firmness, tenderness and flavor of the meat of a small subset of samples. However, the trained judges perceived the meat from highly protein-efficient pigs as less juicy than from pigs that had average or below-average PE, but to investigate if this would affect the consumers’ buying decision, a test with a consumer panel should be conducted.

The genetic correlations of FCR also show possible unfavourable relationships with meat lightness, meat redness, meat yellowness, IMF and cooking loss, which suggest that an indirect selection for higher PE by selecting for lower FCR in Swiss Large White pigs would not only result in lower genetic gain due to the moderate genetic correlation between PE and FCR, but also lead to lighter meat. This agrees with the study of Saintilan et al. [12] who reported significant negative genetic correlations between FCR and meat lightness for Large White dam and sire breed.

## Conclusions

Our results clearly show that PE is heritable, and breeding for protein-efficient pigs is possible. It should be noted that the genetic parameters of PE were estimated here predominantly under conditions of low dietary protein availability. Together with the phenotypic results of the influence of diet presented in this work, and the still not resolved question of genotype-by-diet interactions, this shows the importance of the dietary environment, and the influence of dietary protein content should be considered accordingly. There seems to be a potential for indirectly selecting improved phosphorus efficiency due to its high genetic correlation with PE. Selection for PE does not appear to have major conflicts with meat quality and carcass traits, except for meat redness and yellowness, IMF and cooking loss, which may need to be closely monitored. As for production traits, genetic correlations are favourable, though average daily gain should be monitored to avoid slower growth with increased PE. To achieve significant reduction in pollutant levels in manure, selection on conventional efficiency traits, such as FCR or RFI alone, might not be sufficient. Furthermore, we show that FCR showed genetic antagonism with meat quality traits in the population studied. Thus, including PE in the breeding goals would better contribute to a sustainable and environmentally friendly pig production.

## Declarations

### Ethics approval and consent to participate

The experimental procedure was approved by the Office for Food Safety and Veterinary Affairs (2018_30_FR) and all procedures were conducted in accordance with the Ordinance on Animal Protection and the Ordinance on Animal Experimentation.

### Consent for publication

Not applicable

### Availability of data and materials

The data that support the findings of this study and the code used for models and the statistical analysis are publicly available in Zenodo (DOI:10.5281/zenodo.6985500)

### Competing interests

The authors report no conflicts of interest with any of the data presented.

### Funding

This research was supported by the Fondation Sur-la-Croix to G.B. and C.K.

### Authors’ contributions

EOE carried out the experiment, curated and analyzed the data and drafted the manuscript. GB participated in the design and coordination of the study and in the data collection. CK conceived of the study and participated in its design and coordination. All authors read and approved the final manuscript.

## Acknowledgements

We are grateful to Guy Maïkoff and his team for the maintenance and slaughter of the pigs, for feed production and assistance with DXA scans and Marion Girard for the formulation of the protein-reduced diet. Patrick Schlegel also provided help with the DXA scans. Sébastien Dubois and Paolo Silacci and their teams conducted the chemical analyses of feed and meat and helped with the collection of technological meat quality data, for which Martin Scheeder from Suisag shared protocols with us. Our thanks also go to Barbara Guggenbühl and her team for carrying out the sensory analysis, and Markus Neuditschko for comments on the manuscript.

## Additional files

**Additional file 2 Figure S1-S2, Table S1-S2**

Format: DOC

Title: Figure S1:

Description: Posterior distributions for nutrient efficiencies and production traits.

Title: Figure S2:

Description: Figures to illustrate the overlap of the posterior distributions of meat and carcass quality traits with null distributions.

Title: Table S1:

Description: Table of heritability estimates from MCMCglmm and ASReml-R with or without the inclusion of litter effect

Title: Table S2:

Description: table of heritability, genetic correlations and phenotypic correlation of dressing percentage with protein efficiency

## Footnotes

1 Fütterungsempfehlungen und Nährwerttabellen für Schweine (Feeding recommendations and nutrient tables for pigs). Agroscope, Posieux, Switzerland. Retrieved on 31 January 2017 from https://www.agroscope.admin.ch/agroscope/fr/home/services/soutien/aliments-pour-animaux/apports-alimentaires-recommandes-pour-les-porcs.html

2 Bio-Suisse: Production standards (https://international.bio-suisse.ch/en/import-with-bio-suisse/documents-and-downloads.html), retrieved on March 15, 2023

## Notes

### Competing Interest Statement

The authors have declared no competing interest.

### Summary of Updates

We revised the manuscript following the suggestions of two reviewers. In addition, we switched the analysis of genetic parameters from MCMCglmm to ASReml-R. During the preparation of the revised version after the editor's comments, we reran the animal models to obtain heritability estimates and genetic correlations that were previously done in MCMCglmm in ASReml-R. For most traits in the manuscript, the estimates were very similar between the software packages. However, for protein efficiency, the core trait of this manuscript, the estimates were markedly different (h^2^=0.43 in MCMCglmm vs h^2^=0.60 in ASReml-R). It seemed that MCMCglmm attributes part of the variation to the additive genetic variance components, and part of it to the litter variance component, while ASReml-R attributed most variation to the additive genetic variance. This continued to puzzle us, and we tried to follow this up as good as possible. Thus, to ascertain how accurately MCMCglmm analyses our data, we simulated several data sets using information from previous heritability estimations of protein efficiency with differing litter effects (0.01 - 0.20) but the same heritability (i.e., h^2^ = 0.43). With these data, we ran animal models in both ASReml and MCMCglmm (and brms, another Bayesian mixed-effects model software package). We realized that, while ASReml gave litter effect estimates that were close to the simulated data, MCMCglmm overestimated the litter effect, thereby reducing the heritability. We assume that using the default uniformative prior in MCMCglmm for both effects, additive genetics and litter, resulted in the overestimation of the litter effect at the expense of the additive genetic effect. However, the use of a combination of an uninformative (for the random animal effect) and a parameter-expanded (for the random litter effect) prior gave litter effect estimates close to the true values in the simulated data, but we were not certain if it is proper to use such a combination of priors. Thus, we switched our analysis from MCMCglmm to ASReml. Moreover, the default prior in brms also yielded a result very close to the one of ASReml-R. To allow some degree of comparability with our previous article in Journal of Animal Breeding and Genetics (https://doi.org/10.1111/jbg.12472), we report the results of the MCMCglmm analysis in the Supplementary Material.

