## Supplemental Figures and Tables for "Genetic analysis of protein efficiency and its association with performance and meat quality traits under a protein-restricted diet"

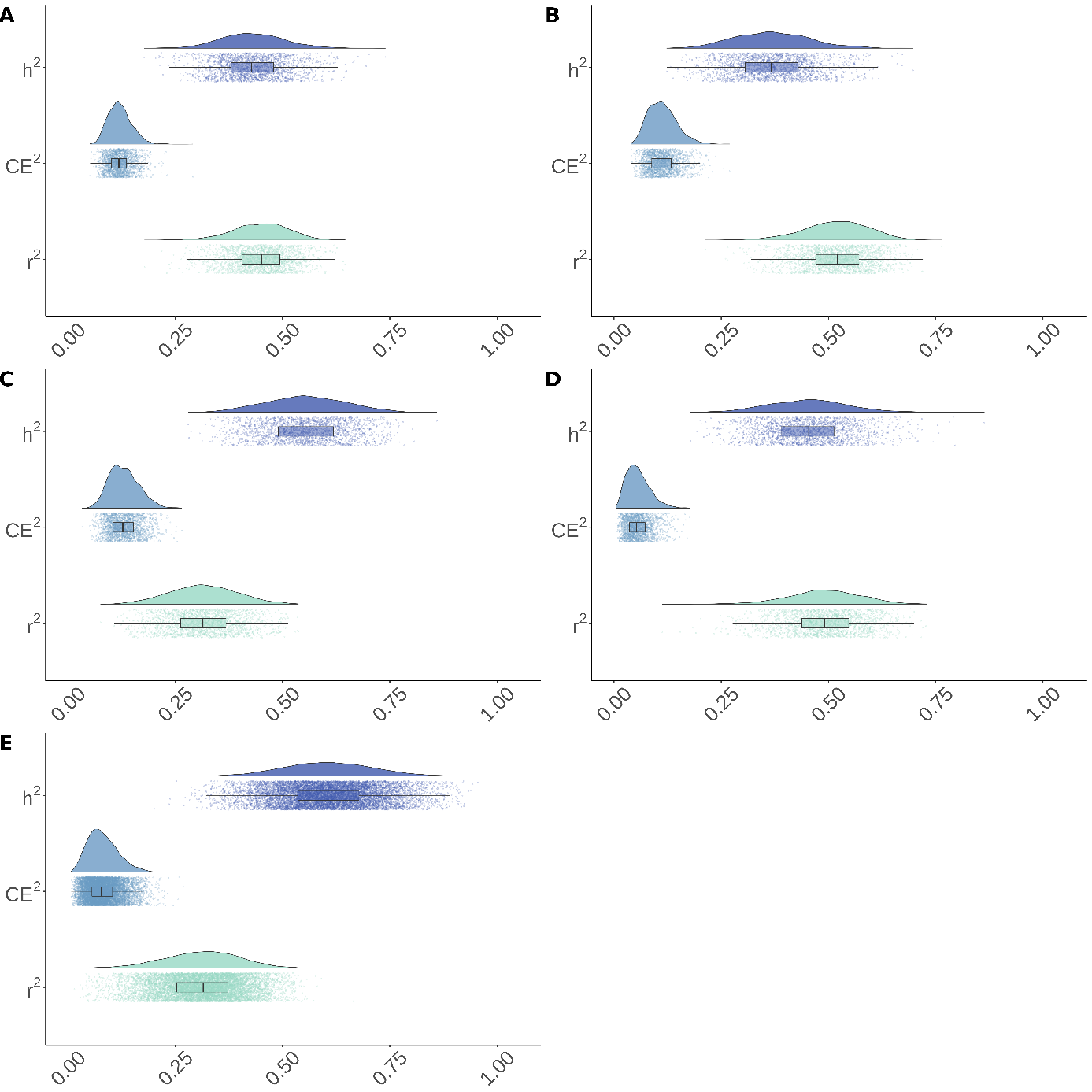

**Figure S1:** Heritability (h^2^), common environment effect (CE^2^; the ratio of the variance of litter to the phenotypic variance) and residual variance (r^2^; the ratio of the residual variance to the phenotypic variance) of protein efficiency (A), phosphorus efficiency (B), average daily gain (C), feed conversion ratio (D), and average daily feed intake (E). Posterior distributions of the respective variance components (upper part), points representing single estimates are shown together with a box plot (with median; whiskers represent the 95% credible interval).

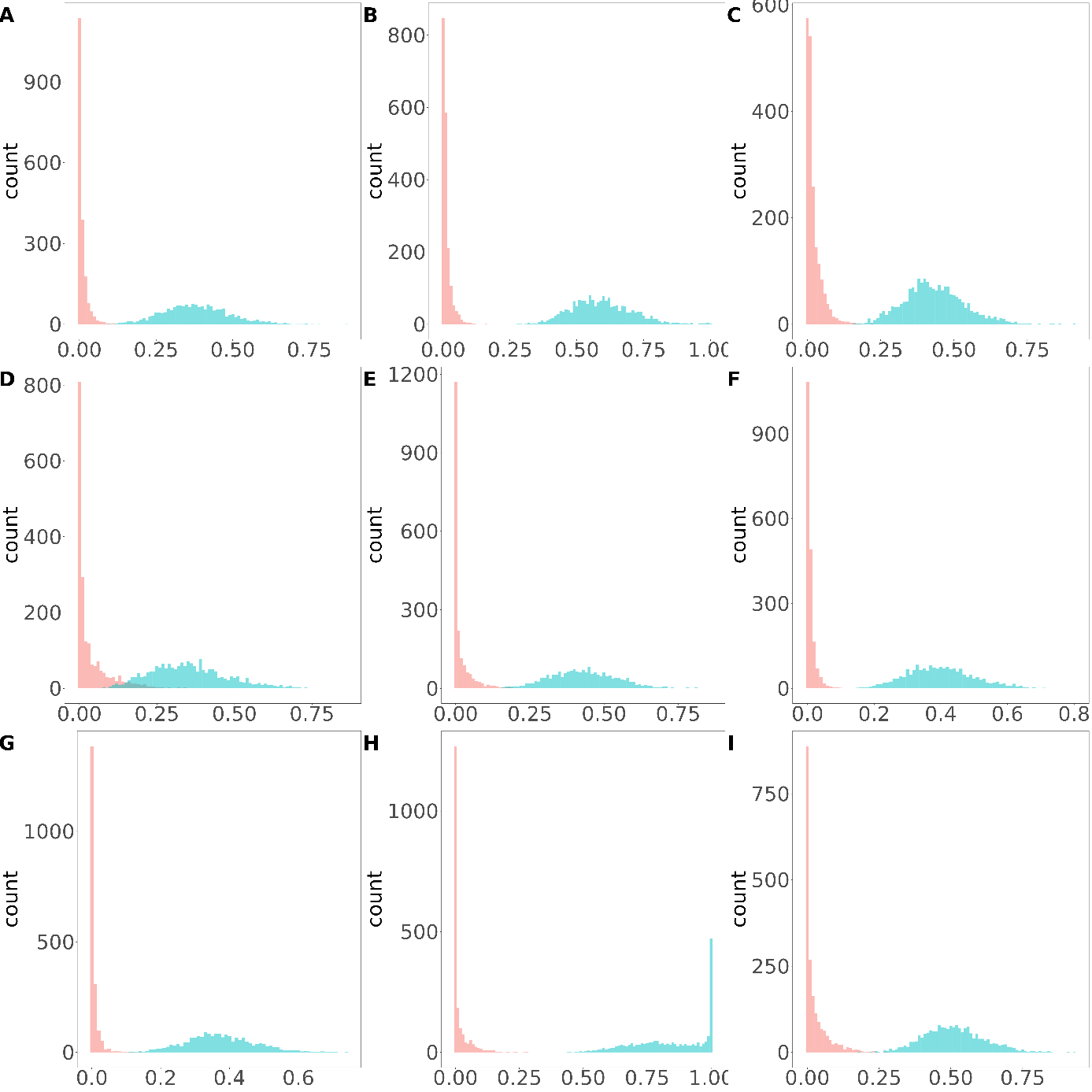

**Figure S2:** Overlap of real posterior distributions of heritability of meat and carcass quality traits with a random distribution. **A** meat lightness, **B** meat redness, **C** meat yellowness, **D** shear force, **E** drip loss, **F** cooking loss, **G** intramuscular fat content, **H** backfat thickness.

**Table S1:** Heritability estimates from ASREML-R and MCMCglmm with and without the random litter effect

|  | ASReml | | MCMCglmm | |
| --- | --- | --- | --- | --- |
| Trait | Heritability with random litter effect | Heritability without random litter effect | Heritability with random litter effect | Heritability without random litter effect |
| Protein efficiency | 0.60 ± 0.08 | 0.60 ± 0.08 | 0.43 (0.29 – 0.58) | 0.60 (0.49 – 0.76) |
| Phosphorus efficiency | 0.46 ± 0.11 | 0.54 ± 0.08 | 0.36 (0.19 – 0.54) | 0.49 (0.40 – 0.71) |
| ADG | 0.59 ± 0.11 | 0.73 ± 0.08 | 0.54 (0.39 – 0.75) | 0.70 (0.59 – 0.91) |
| FCR | 0.47 ± 0.10 | 0.52 ± 0.08 | 0.46 (0.27 – 0.63) | 0.54 (0.38 – 0.73) |
| ADFI | 0.59 ± 0.11 | 0.73 ± 0.08 | 0.61 (0.41 – 0.83) | 0.76 (0.60 – 0.92) |
| IMF | - | 0.73 ± 0.12 | - | 0.99 (0.58 – 0.99) |
| LMA | - | 0.34 ± 0.11 | - | 0.31 (0.12 – 0.61) |
| L colour | - | 0.37 ± 0.10 | - | 0.32 (0.17 – 0.60) |
| a colour | - | 0.57 ± 0.11 | - | 0.58 (0.37 – 0.83) |
| b colour | - | 0.41 ± 0.10 | - | 0.37 (0.22 – 0.62) |
| Shear force | - | 0.42 ± 0.10 | - | 0.43 (0.21 – 0.64) |
| Drip loss | - | 0.39 ± 0.09 | - | 0.37 (0.21 – 0.59) |
| Cooking loss | - | 0.36 ± 0.09 | - | 0.38 (0.18 – 0.56) |
| Backfat thickness | - | 0.50 ± 0.10 | - | 0.48 (0.30 – 0.72) |

**Table S2:** Genetic correlations (above diagonal) and phenotypic correlation (lower diagonal) of dressing percentage with protein efficiency

|  | Protein efficiency | Warm dressing percentage | Cold dressing percentage |
| --- | --- | --- | --- |
| Protein efficiency |  | 0.28 ± 0.19 | 0.31 ± 0.18 |
| Warm dressing percentage | 0.14 ± 0.05 |  | 0.23 ± 0.13 |
| Cold dressing percentage | 0.16 ± 0.05 | 0.32 ± 0.13 |  |
